# Comparative analyses of tailbeat frequency and stride length reveal how regionally endothermic fishes cruise fast

**DOI:** 10.64898/2026.08.14.744074

**Authors:** Soma Tokunaga, Nicholas L. Payne, Ryo Kawabe, Itsumi Nakamura, Seishiro Furukawa, Wei-Chuan Chiang, Jayson M. Semmens, Carl G. Meyer, Yuuki Y. Watanabe

**Affiliations:** Department of Evolutionary Studies of Biosystems, The Graduate University for Advanced Studies, SOKENDAI, Hayama, Kanagawa, Japan; Hawai‘i Institute of Marine Biology, University of Hawai‘i at Mānoa, Kāne‘ohe, HI, USA; School of Natural Sciences, Department of Zoology, Trinity College Dublin, Dublin, Ireland; Institute for East China Sea Research, Organization for Marine Science and Technology, Nagasaki University, Nagasaki, Japan; Japan Fisheries Research and Education Agency, Yokohama, Kanagawa, Japan; Eastern Fishery Research Center, Fisheries Research Institute, Ministry of Agriculture, Chenggong, Taiwan; Institute for Marine and Antarctic Studies, University of Tasmania, Hobart, TAS, Australia; Research Center for Integrative Evolutionary Science, The Graduate University for Advanced Studies, SOKENDAI, Hayama, Kanagawa, Japan

**Keywords:** biologging, fish endothermy, lamnid shark, swimming kinematics, tuna

## Abstract

Cruising speed is a key factor affecting prey-search efficiency and migration range in continuously swimming animals. Tunas and lamnid sharks (e.g., white sharks) have convergently evolved traits for high-speed cruising, including the ability to maintain slow-twitch, aerobic red muscle (RM) warmer than ambient water, known as RM endothermy. Despite their well-known high cruising speeds, kinematic features underlying their elevated speeds remain unclear. Swim speed is the product of tailbeat frequency (TBF; Hz) and stride length (SL, the absolute distance traveled per tailbeat; m). RM endothermy is expected to elevate TBF by enhancing muscle contraction performance. Furthermore, within RM-endothermic fishes, tunas and lamnid sharks may exhibit distinct kinematic features because of differences in caudal fin morphology and tailbeat amplitude. Here, we compiled kinematic parameters from 20 fish species, including five RM-endothermic species, measured in the wild using animal-borne sensors. Comparative analyses showed that, for a given body mass and water temperature, RM-endothermic fishes exhibited 1.9 times higher cruising speed and TBF than ectothermic fishes, while SL remained similar. Within RM-endothermic fishes, tunas exhibited 2.3 times higher TBF than similar-sized lamnid sharks, whereas lamnid sharks showed 1.7 times longer SL than similar-sized tunas. These results indicate that RM endothermy is generally associated with higher TBF, while significant kinematic differences remain between tunas and lamnid sharks. This divergence may be partly explained by the greater caudal fin area and tailbeat amplitude in lamnid sharks. It may also reflect contrasting skeletal types of teleosts and elasmobranchs, which potentially influence body stiffness and swimming kinematics.

## INTRODUCTION

Cruising speed is a key factor affecting prey-search efficiency, migration range, and ultimately fitness in continuously swimming animals. Among fishes, tunas and lamnid sharks (e.g., white sharks *Carcharodon carcharias*) are widely recognized as cruising specialists that search large areas for prey, and some exhibit extensive annual migrations of ∼10,000 km (Bernal et al., 2001; Bonfil et al., 2005; Watanabe et al., 2015; Webb, 1984). Despite the deep evolutionary divergence between teleosts and elasmobranchs (∼450 million years ago; Ravi and Venkatesh, 2008), tunas and lamnid sharks share several traits associated with high-speed cruising, including streamlined bodies, lunate caudal fins with high aspect ratio, and a ‘thunniform’ swimming mode characterized by lateral body movement largely restricted to the caudal region (Bernal et al., 2001; Donley et al., 2004).

Another remarkable trait shared by tunas and lamnid sharks is red muscle (RM) endothermy. Unlike most fishes, whose body temperatures are determined by ambient water temperature (i.e., ectothermy), RM-endothermic fishes maintain slow-twitch, aerobic swimming muscles warmer than the ambient water by conserving the generated metabolic heat via vascular countercurrent heat exchangers (Carey and Teal, 1966; Carey and Teal, 1969). Elevated RM temperature may enhance muscle contractile performance, potentially increasing the capacity for high-speed swimming (Carey et al., 1971; Dickson and Graham, 2004). In addition, the elevated metabolic rate associated with RM endothermy (Payne et al., 2026) may shift the swim speed that minimizes cost of transport to a higher value (Claireaux et al., 2006; Whitney et al., 2016). Consistent with these scenarios, previous comparative studies showed that RM-endothermic species exhibit significantly faster cruising speeds and longer migration distances than ectothermic species (Harding et al., 2021; Watanabe et al., 2015).

Despite substantial research on RM-endothermic fishes, kinematic features underlying their elevated cruising speeds remain poorly understood. Swim speed (m/s) is the product of tailbeat frequency (TBF; Hz) and stride length (SL, defined here as the absolute distance traveled per tailbeat; m) (Videler and Wardle, 1991). However, the relative contributions of these factors to high-speed cruising in RM-endothermic fishes remain unclear. This is likely because large pelagic tunas and lamnid sharks are difficult to maintain in captivity, limiting opportunities for detailed kinematic measurements. Although biologging techniques with animal-borne electronic devices would be powerful tools to measure swimming kinematics in the wild (Payne et al., 2014; Watanabe and Papastamatiou, 2023), many studies rely primarily on accelerometers, which can quantify TBF but not swim speed. Because calculating SL requires simultaneous measurements of swim speed and TBF from the same individual, datasets containing both variables are needed to evaluate the relative contributions of TBF and SL to elevated cruising speed.

RM-endothermic fishes may achieve high-speed cruising primarily through increased TBF rather than extended SL. TBF generally decreases with body size and increases with water (or body) temperature (Videler and Wardle, 1991; Watanabe et al., 2012). By contrast, SL is strongly influenced by body size (Bainbridge, 1958; Hunter and Zweifel, 1971; Videler and Wardle, 1991). Temperature effects on SL have been reported, but appear to be largely species-specific (Riyanto et al., 2014; Sisson and Sidell, 1987; Stevens, 1979). In addition, the thunniform swimming mode shared by RM-endothermic fishes may contribute to high TBF and short SL through its limited lateral body movement (Donley and Dickson, 2000; Dowis et al., 2003). The influence of water viscosity is negligible in large fishes that swim at high Reynolds numbers (Webb, 1988). Taken together, we hypothesize that RM-endothermic fishes, which maintain elevated RM temperatures, show higher TBF than ectothermic fishes for a given body size and water temperature.

Within RM-endothermic fishes, however, tunas and lamnid sharks may exhibit significant differences in TBF and SL. Despite their overall convergence for high-speed cruising, marked differences remain in their morphology and swimming kinematics. Morphologically, tunas have apparently smaller caudal fin areas than lamnid sharks (Fig. 1A), although quantitative comparisons are lacking. Kinematically, despite the convergent thunniform swimming mode, tailbeat amplitude (i.e., the peak-to-peak lateral excursion of the caudal fin tip; Fig. 1B) differs between them. Captive studies reported tailbeat amplitudes of 16% of fork length (FL) in Atlantic bluefin tuna *Thunnus thynnus* (Wardle et al., 1989) and 26% of total length (TL) in shortfin mako sharks *Isurus oxyrinchu*s (Donley et al., 2005). Because both greater caudal fin area and tailbeat amplitude contribute to greater thrust generation per tailbeat (Floryan et al., 2018), tunas may compensate for their smaller caudal fin area and tailbeat amplitude by beating their tails more frequently than lamnid sharks.

**Figure 1.**
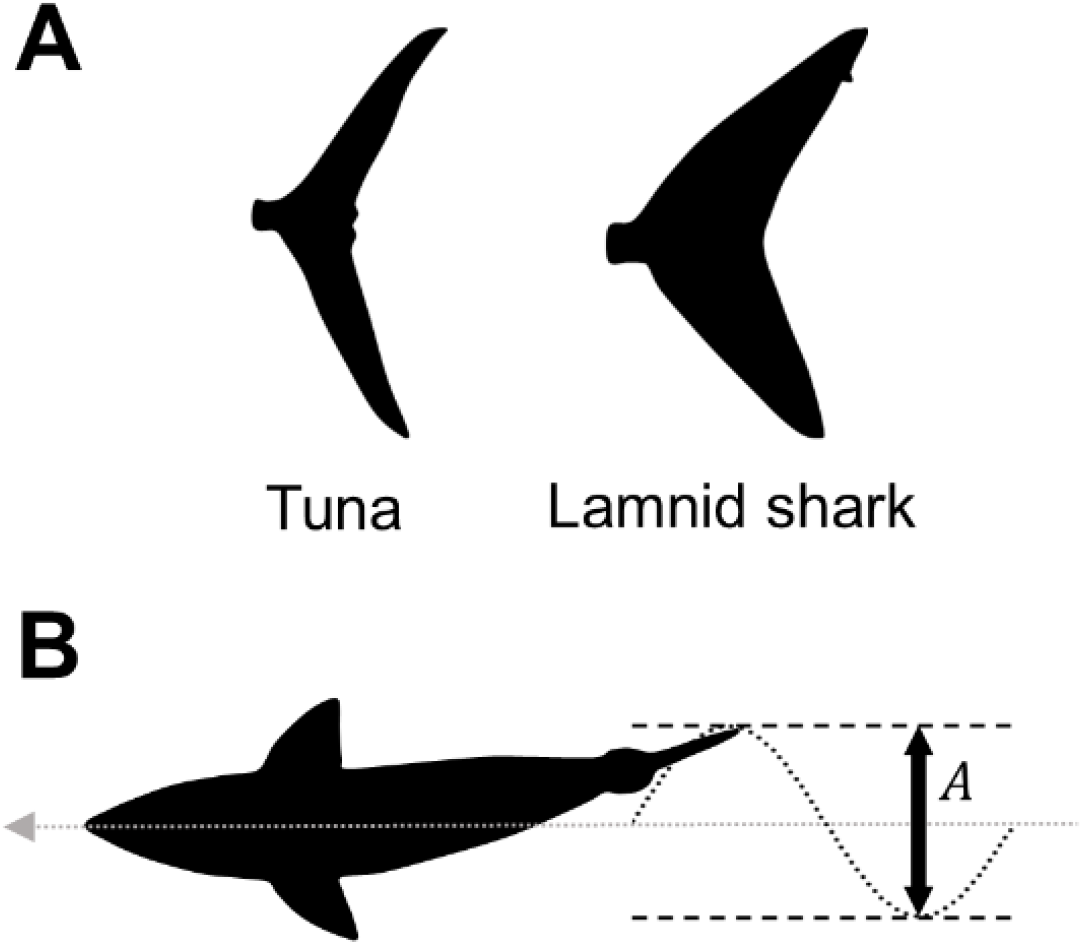
Morphological and kinematic features of tunas and lamnid sharks. (A) Comparison of caudal fin morphology, showing an apparently smaller caudal fin area in tunas than in lamnid sharks. (B) Tailbeat amplitude, *A*, measured as the peak-to-peak lateral excursion of the caudal fin tip. The gray arrow and black dotted line indicate the swimming direction and the trajectory of the tail tip, respectively.

In this study, we examined kinematic features underlying high-speed cruising in RM-endothermic fishes. By compiling original biologging datasets from a variety of continuously swimming fishes in the wild, we conducted comparative analyses to test the following two hypotheses: (1) RM-endothermic fishes exhibit significantly higher TBF than ectothermic fishes for a given size and water temperature, and (2) within RM-endothermic fishes, tunas show significantly higher TBF and shorter SL than similar-sized lamnid sharks.

## MATERIALS AND METHODS

### Data compilation

To examine swimming behavior under natural conditions, original datasets obtained from field experiments were compiled from both published and unpublished sources. Swim speed (m/s) has been measured primarily using two approaches: (1) direct measurement using animal-borne propeller speed sensors and (2) indirect estimation from acoustic tracking and reconstructed movement paths (Watanabe et al., 2015). Because the latter can be influenced by ocean currents, only data obtained using propeller speed sensors, which directly measure swim speed relative to water, were included. To minimize methodological bias, analyses were restricted to deployments of PD2GT, PD2GTL, PD3GT, or 3MPD3GT loggers (all manufactured by Little Leonardo Corp., Tokyo, Japan), which recorded bi- or tri-axial acceleration (g), depth (m), and ambient water temperature (°C) simultaneously.

Unpublished data were compiled from field experiments conducted on the following species: a bigeye thresher shark *Alopias superciliosus* (May–June 2016, off Taitung, southeastern Taiwan), a blacktip shark *Carcharhinus limbatus* (March 2014, inside Kāneʻohe Bay, Hawaiʻi, USA), broadnose sevengill sharks *Notorynchus cepedianus* (April 2014, off Tasmania, Australia), a striped marlin *Kajikia audax* (November 2014, off Ōarai, Ibaraki, Japan), a greater amberjack *Seriola dumerili* (September 2021, off Tanegashima Island, Kagoshima, Japan), and a yellowtail amberjack *Seriola lalandi* (August 2022, off Fukue Island, Nagasaki, Japan). Animals were hooked and restrained alongside a boat or brought onboard for tag deployment. The tagging package consisted of a PD2GT (21 mm in diameter, 117 mm in length, 60 g), PD2GTL (24 mm in diameter, 124 mm in length, 64 g), PD3GT (21 mm in diameter, 115 mm in length, 60 g), or 3MPD3GT logger (16.5 mm in diameter, 83.5 mm in length, 42 g) (Little Leonardo Corp.). The package also included a float, a time-scheduled release mechanism (Little Leonardo), a VHF radio transmitter (Advanced Telemetry Systems), and a satellite transmitter (Wildlife Computers). The package was attached to the body surface, except for the striped marlin, for which the package was mounted on the upper bill. After 1–4 days of recording, the package detached, floated, was located using VHF and satellite signals, and was recovered by boat. The logger recorded propeller rotations at 0.5 or 1 s intervals, depth and temperature at 1 s intervals, and bi- or tri-axial acceleration at 1/16, 1/20, 1/32, or 1/50 s intervals. Propeller rotations were converted to swim speed (m/s) using calibration equations (Watanabe et al., 2008). The attachment angle of the data logger, defined as the difference between the pitch angles of the fish body and the logger, was estimated following Kawatsu et al. (2009). Swim speed was then calibrated by dividing the speed by the cosine of the attachment angle, following Watanabe et al. (2019b).

Body mass, rather than body length, was used as the index of body size because our dataset included morphologically diverse fish taxa. Body mass (kg) was estimated from total length (TL), fork length (FL), or precaudal length (PCL) using species-specific length–weight relationships derived from the literature or FishBase (Tables S1, S2), except for the striped marlin, for which body mass was estimated at capture. The presence or absence of RM endothermy for each species was determined based on previous studies (Dickson and Graham, 2004; Sepulveda et al., 2005). Body temperatures of Atlantic bluefin tuna were estimated from a relationship with ambient water temperature (Addis et al., 2009). Body temperatures of Pacific bluefin tuna *Thunnus orientalis* were estimated from a relationship between temperature difference (i.e., body temperature – water temperature) and body mass (Kitagawa et al., 2006). Body temperatures of white sharks and a salmon shark *Lamna ditropis* were estimated from mean body temperatures across individuals under similar thermal conditions (Goldman, 1997; Goldman et al., 2004). Body temperatures of shortfin mako sharks were directly measured with stalk temperature sensors deployed on free-swimming individuals (Tokunaga et al., 2025). Body temperatures of ectothermic fishes were assumed to equal ambient water temperature.

### Data analyses

Data were analyzed using Igor Pro (WaveMetrics Inc.) with the Ethographer package (Sakamoto et al., 2009). To remove potential effects of capture, the first 6 h of each deployment were excluded (Iosilevskii et al., 2022). White sharks were an exception because they were tagged without capture (Watanabe et al., 2019b). In this case, the initial 6 h were retained, but periods during which sharks interacted with cage diving activity were excluded (Huveneers et al., 2018). Periods of apparent propeller malfunction, defined as prolonged periods of near-zero or unusually low swim speed despite continued swimming activity in the acceleration data, were also excluded.

TBF (Hz) was calculated at 1 s intervals by spectral analysis of lateral acceleration using the peak tracer function in Ethographer (Nakamura et al., 2011; Sakamoto et al., 2009). Swim speed, TBF, and depth were subsampled at 1 min intervals, and these data were separated into horizontal swimming, ascent, or descent based on vertical speed. Only the horizontal swimming with vertical speeds between −0.1 and +0.1 m/s was used to exclude the effects of buoyancy-driven behaviors such as gliding (Nakamura et al., 2011; Watanabe et al., 2019b). Mean values of swim speed (i.e., cruising speed), TBF, and water and body temperature during horizontal swimming were calculated for each individual, and SL (m) was calculated as mean swim speed divided by mean TBF. SL was therefore a derived rather than directly measured kinematic variable.

To compare RM-endothermic and ectothermic fishes, cruising speed, TBF, and SL were analyzed separately as response variables. Body mass and temperature (water or body) were included as continuous predictors, and RM endothermy (presence or absence) was included as a categorical predictor. Water and body temperature were modeled separately to avoid multicollinearity. Cruising speed, TBF, SL, and body mass were log_10_-transformed to improve linearity. To account for phylogenetic non-independence among species (Felsenstein, 1985), Bayesian generalized linear mixed models were fitted using the MCMCglmm package in R (Hadfield, 2010), with phylogeny and species identity modeled as phylogenetic and non-phylogenetic random effects, respectively. A phylogenetic tree was constructed in the software Mesquite (Maddison and Maddison, 2019) based on the Chondrichthyan Tree of Life (Naylor, 2026) for elasmobranchs and the Fish Tree of Life (Rabosky et al., 2018) for teleosts, with branch lengths scaled by Grafen’s method (Grafen, 1989) (Fig. S1). Using weakly informative priors (V = 1, nu = 0.002), each model was run for 1,000,000 iterations, with the first 100,000 discarded as burn-in and every 500th iteration retained for posterior sampling, yielding effective sample sizes of >1000 for all parameters. Predictor effects were considered statistically significant when their 95% credible intervals excluded zero. To obtain the final model, predictors whose 95% credible intervals overlapped zero were sequentially removed from the full model. When multiple predictors met this criterion, the predictor with the smallest absolute posterior mean was removed first.

To compare RM-endothermic tunas and lamnid sharks, the same set of continuous predictors was used, but clade (teleost vs. elasmobranch, that is, tuna vs. lamnid shark) was included as a categorical predictor instead of RM endothermy. Because teleosts and elasmobranchs diverged early in fish evolution (∼450 million years ago; Ravi and Venkatesh, 2008), clade itself represents a strong phylogenetic contrast. The effects of clade and phylogenetic relatedness could therefore not be reliably distinguished if both were included in the same model. Accordingly, these analyses included species identity as a non-phylogenetic random effect but did not include phylogeny as a phylogenetic random effect. Model selection followed the same procedure described above.

## RESULTS

Kinematic parameters were obtained from 66 free-ranging individuals representing 20 fish species, including five species with RM endothermy. Estimated body mass ranged from 3.9 to 3250 kg, water temperature ranged from 0.3 to 29.0°C, and body temperature ranged from 0.3 to 30.9°C (Tables 1, S3).

Cruising speed was significantly affected by temperature (water or body temperature) and RM endothermy, but not by body mass (Table 2; Fig. 2A). For a given water temperature, cruising speed was 1.9 times faster in RM-endothermic fishes (speed = 0.53 × 10^0.014Temp^) than in ectothermic fishes (speed = 0.28 × 10^0.014Temp^) (Fig. 3A). For a given body temperature, cruising speed was 1.4 times faster in RM-endothermic fishes (speed = 0.34 × 10^0.017Temp^) than in ectothermic fishes (speed = 0.24 × 10^0.017Temp^) (Fig. 3B). However, in the full model including body mass and body temperature, the effect of RM endothermy was not significant (i.e., the 95% credible interval included zero) (Table S4). Moreover, after removing whale sharks with exceptionally large body masses (708–3250 kg) and Greenland sharks with exceptionally low body temperatures (0.3–1.0°C), the final model included body mass and body temperature but did not include RM endothermy (Table S5).

**Figure 2.**
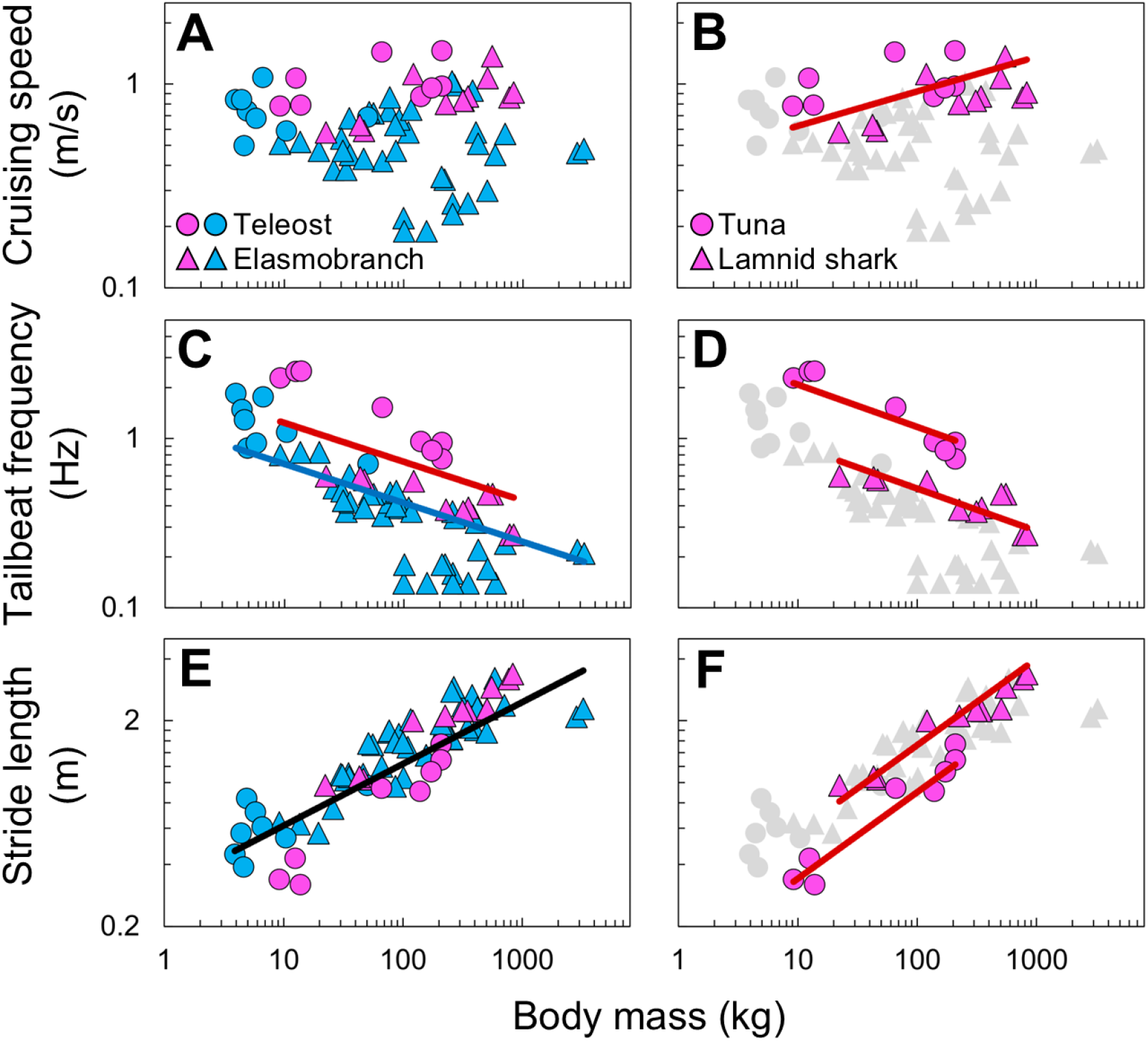
Relationships between kinematic parameters and body mass in fishes. Cruising speed (A, B), tailbeat frequency (C, D), and stride length (E, F) are plotted against body mass. Each point represents an individual; circles and triangles indicate teleosts and elasmobranchs, respectively. Panels A, C, and E compare RM-endothermic (pink) and ectothermic (sky blue) fishes. Panels B, D, and F compare RM-endothermic tunas (Atlantic and Pacific bluefin tunas) and lamnid sharks (white, salmon, and shortfin mako sharks), with ectothermic fishes shown in gray for visual reference. Regression lines are shown in red for RM-endothermic fishes, blue for ectothermic fishes, and black when there was no significant difference between the two groups. No regression line is shown when the effect of body mass is not significant. Regression equations are provided in the main text.

**Figure 3.**
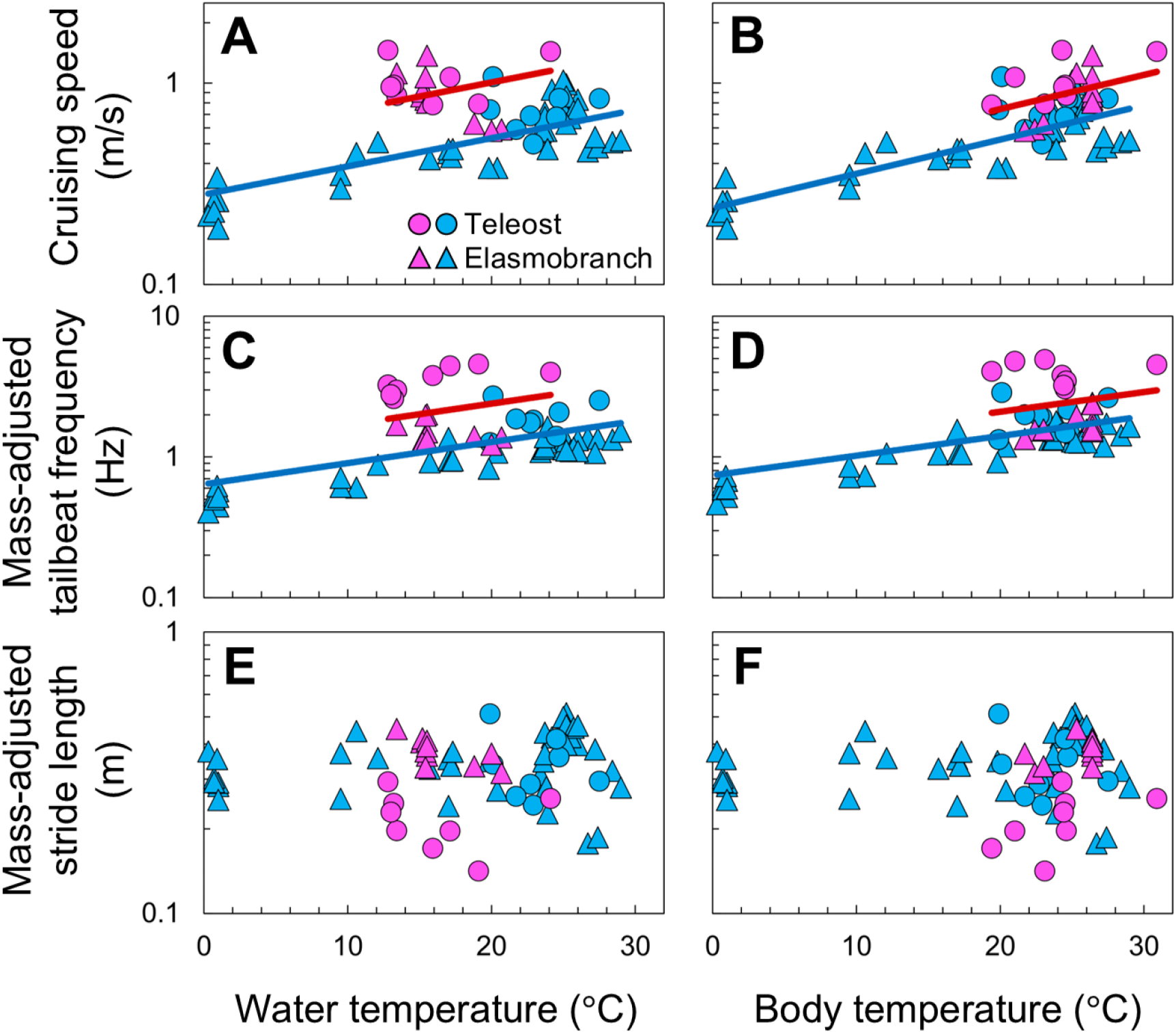
Relationships between kinematic parameters and temperature in fishes. Cruising speed (A, B), mass-adjusted tailbeat frequency (C, D), and mass-adjusted stride length (E, F) are plotted against water temperature (A, C, E) and body temperature (B, D, F). Cruising speed is shown without mass adjustment because body mass is not retained in the final model. TBF and SL are standardized to a 1-kg body mass using the allometric slopes from the corresponding final models (Table 2). Each point represents an individual; circles and triangles indicate teleosts and elasmobranchs, respectively; pink and sky blue indicate RM-endothermic and ectothermic fishes, respectively. Regression lines are shown in red for RM-endothermic fishes and blue for ectothermic fishes. No regression line is shown when the effect of temperature is not significant. Regression equations are provided in the main text.

**Table 1.** Summary of body size and kinematic parameters for 20 fish species.

| Species | n | Fork length (m)<sup>a</sup> | Body mass (kg)<sup>b</sup> | Cruising speed (m/s) | Tailbeat frequency (TBF, Hz) | Stride length (SL, m) | Water temp (°C) | Body temp (°C) | RM endo-thermy | Reference |
| --- | --- | --- | --- | --- | --- | --- | --- | --- | --- | --- |
| White shark<br>*Carcharodon carcharias* | 7 | 2.68–4.00 | 226–834 | 0.80–1.37 | 0.27–0.47 | 2.11–3.37 | 15.1–15.5 | 26.4 | Yes | Watanabe et al. (2019b) |
| Salmon shark<br>*Lamna ditropis* | 1 | 1.95 | 121 | 1.12 | 0.56 | 2.00 | 13.4 | 25.3 | Yes | Watanabe et al. (2015) |
| Shortfin mako shark<br>*Isurus oxyrinchus* | 3 | 1.29–1.63 | 22.3–46.5 | 0.58–0.63 | 0.57–0.60 | 0.97–1.07 | 18.8–20.7 | 21.7–23.0 | Yes | Tokunaga et al. (2025) |
| Bigeye thresher shark<br>*Alopias superciliosus* | 1 | 1.69 | 66.3 | 0.42 | 0.35 | 1.20 | 15.7 | 15.7 | No | This study |
| Whale shark<br>*Rhincodon typus* | 3 | 4.00–6.58 | 708–3250 | 0.46–0.57 | 0.21–0.24 | 2.09–2.38 | 23.4–27.4 | 23.4–27.4 | No | Nakamura et al. (2020) |
| Blue shark<br>*Prionace glauca* | 2 | 1.61–1.74 | 25.9–33.0 | 0.38 | 0.37–0.51 | 0.75–1.03 | 19.8–20.4 | 19.8–20.4 | No | Watanabe et al. (2021) |
| Tiger shark<br>*Galeocerdo cuvier* | 10 | 1.55–3.28 | 35.0–403 | 0.58–1.03 | 0.32–0.61 | 1.11–2.91 | 23.6–26.0 | 23.6–26.0 | No | Nakamura et al. (2011); Tokunaga et al. (2025) |
| Blacktip shark<br>*Carcharhinus limbatus* | 1 | 1.18 | 19.5 | 0.47 | 0.83 | 0.57 | 23.9 | 23.9 | No | This study |
| Blacktip reef shark<br>*Carcharhinus melanopterus* | 2 | 0.93–1.05 | 9.2–13.6 | 0.51–0.52 | 0.80–0.83 | 0.63–0.64 | 28.4–29.0 | 28.4–29.0 | No | Watanabe et al. (2015) |
| Gray reef shark<br>*Carcharhinus amblyrhynchos* | 1 | 1.29 | 30.0 | 0.54 | 0.49 | 1.10 | 27.2 | 27.2 | No | Watanabe et al. (2015) |
| Oceanic whitetip shark<br>*Carcharhinus longimanus* | 4 | 1.74–2.35 | 51.4–116 | 0.63–0.75 | 0.37–0.47 | 1.55–2.03 | 25.2–26.0 | 25.2–26.0 | No | Watanabe et al. (2015) |
| Greenland shark<br>*Somniosus microcephalus* | 7 | 2.06–3.10 | 100–346 | 0.19–0.34 | 0.14–0.18 | 1.06–1.89 | 0.3–1.0 | 0.3–1.0 | No | Watanabe et al. (2012) |
| Bluntnose sixgill shark<br>*Hexanchus griseus* | 4 | 2.83–4.05 | 208–587 | 0.30–0.51 | 0.14–0.22 | 1.76–3.21 | 9.5–12.1 | 9.5–12.1 | No | Nakamura et al. (2015) |
| Broadnose sevengill shark<br>*Notorynchus cepedianus* | 4 | 1.46–1.93 | 31.0–86.4 | 0.43–0.47 | 0.39–0.49 | 0.96–1.10 | 17.0–17.3 | 17.0–17.3 | No | This study |
| Atlantic bluefin tuna<br>*Thunnus thynnus* | 4 | 1.90–2.18 | 139–209 | 0.87–1.46 | 0.76–0.96 | 0.91–1.54 | 12.8–13.4 | 24.3–24.6 | Yes | Harding et al. (2021) |
| Pacific bluefin tuna<br>*Thunnus orientalis* | 4 | 0.77–1.47 | 9.2–65.8 | 0.78–1.44 | 1.53–2.50 | 0.32–0.94 | 15.9–24.1 | 19.4–30.9 | Yes | Furukawa et al. (2014) |
| Common dolphinfish<br>*Coryphaena hippurus* | 5 | 0.79–0.90 | 3.9–5.8 | 0.50–0.84 | 0.88–1.85 | 0.39–0.84 | 19.9–27.5 | 19.9–27.5 | No | Furukawa et al. (2011) |
| Striped marlin<br>*Kajikia audax* | 1 | 1.68 | 50 | 0.69 | 0.71 | 0.97 | 22.7 | 22.7 | No | This study |
| Greater amberjack<br>*Seriola dumerili* | 1 | 0.81 | 6.6 | 1.08 | 1.76 | 0.61 | 20.1 | 20.1 | No | This study |
| Yellowtail amberjack<br>*Seriola lalandi* | 1 | 0.95 | 10.4 | 0.59 | 1.09 | 0.54 | 21.7 | 21.7 | No | This study |
| Total | 66 | 0.77–6.58 | 3.9–3250 | 0.19–1.46 | 0.14–2.50 | 0.32–3.37 | 0.3–29.0 | 0.3–30.9 |  |  |

<sup>a</sup> Missing FL values were estimated from TL using species-specific TL–FL relationships (Table S1), except for the striped marlin, for which eye-fork length (EFL) was back-calculated from the estimated body mass using the length–weight relationship (Table S2).
<sup>b</sup> Body mass was estimated from species-specific length–weight relationships (Table S2), except for the striped marlin, for which body mass was estimated at capture.

**Table 2.** Posterior parameter estimates for final models from the MCMCglmm analyses.

| Comparison | Model | Predictor | Posterior Mean | 95% credible interval |  |
| --- | --- | --- | --- | --- | --- |
|  |  |  |  | Lower | Upper |
| RM endothermy vs. Ectothermy | $\log_{10}(\text{Speed}) \sim \text{Water temp} + \text{Endothermy}$ | Water temp | 0.014 | 0.0061 | 0.024 |
|  |  | Endothermy | 0.28 | 0.14 | 0.43 |
| | $\log_{10}(\text{Speed}) \sim \text{Body temp} + \text{Endothermy}$ | Body temp | 0.017 | 0.010 | 0.024 |
|  |  | Endothermy | 0.15 | 0.024 | 0.29 |
| | $\log_{10}(\text{TBF}) \sim \log_{10}(\text{Mass}) + \text{Water temp} + \text{Endothermy}$ | $\log_{10}(\text{Mass})$ | -0.23 | -0.29 | -0.18 |
|  |  | Water temp | 0.015 | 0.0081 | 0.023 |
|  |  | Endothermy | 0.29 | 0.14 | 0.44 |
| | $\log_{10}(\text{TBF}) \sim \log_{10}(\text{Mass}) + \text{Body temp} + \text{Endothermy}$ | $\log_{10}(\text{Mass})$ | -0.26 | -0.30 | -0.20 |
|  |  | Body temp | 0.014 | 0.0073 | 0.021 |
|  |  | Endothermy | 0.17 | 0.037 | 0.33 |
| | $\log_{10}(\text{SL}) \sim \log_{10}(\text{Mass})$ | $\log_{10}(\text{Mass})$ | 0.30 | 0.24 | 0.36 |
| Tuna vs. Lamnid shark | $\log_{10}(\text{Speed}) \sim \log_{10}(\text{Mass})$ | $\log_{10}(\text{Mass})$ | 0.17 | 0.0024 | 0.32 |
| | $\log_{10}(\text{TBF}) \sim \log_{10}(\text{Mass}) + \text{Clade}$ | $\log_{10}(\text{Mass})$ | -0.25 | -0.37 | -0.12 |
|  |  | Clade | 0.36 | 0.12 | 0.63 |
| | $\log_{10}(\text{SL}) \sim \log_{10}(\text{Mass}) + \text{Clade}$ | $\log_{10}(\text{Mass})$ | 0.42 | 0.32 | 0.53 |
|  |  | Clade | -0.22 | -0.39 | -0.017 |

Water and body temperature were included separately to avoid multicollinearity. Final models were obtained by sequentially removing predictors with 95% credible intervals overlapping zero. The clade variable indicates whether a species is a tuna or a lamnid shark.

TBF was significantly affected by body mass, temperature (water or body temperature), and RM endothermy (Table 2). For a given mass, TBF was 1.8 times higher in RM-endothermic fishes (TBF = 2.1 × Mass^-0.23^) than in ectothermic fishes (TBF = 1.2 × Mass^-0.23^) (Fig. 2C). For a given mass and water temperature, TBF was 1.9 times higher in RM-endothermic fishes (TBF = 1.2 × Mass^-0.23^ × 10^0.015Temp^) than in ectothermic fishes (TBF = 0.64 × Mass^-0.23^ × 10^0.015Temp^) (Fig. 3C). For a given mass and body temperature, TBF was 1.5 times higher in RM-endothermic fishes (TBF = 1.1 × Mass^-0.26^ × 10^0.014Temp^) than in ectothermic fishes (TBF = 0.74 × Mass^-0.26^ × 10^0.014Temp^) (Fig. 3D). By contrast, SL was significantly affected only by body mass (SL = 0.31 × Mass^0.30^) (Table 2; Figs. 2E, 3E, 3F). The effects of RM endothermy on TBF and SL were largely unchanged in the full models (Table S4) or in the models excluding whale sharks and Greenland sharks (Table S5).

Within RM-endothermic fishes, cruising speed was significantly affected only by body mass (speed = 0.42 × Mass^0.17^), with no significant difference between tunas and lamnid sharks for a given mass (Table 2; Fig. 2B). In contrast, both TBF and SL were significantly affected by body mass and clade (Table 2). For a given mass, TBF was 2.3 times higher in tunas (TBF = 3.7 × Mass^-0.25^) than in lamnid sharks (TBF = 1.6 × Mass^-0.25^) (Figs. 2D, 4), whereas SL was 1.7 times longer in lamnid sharks (SL = 0.22 × Mass^0.42^) than in tunas (SL = 0.13 × Mass^0.42^) (Fig. 2F). Across all response variables, the statistical significance of the clade effect was consistent between the final models and their corresponding full models (Table S4).

**Figure 4.**
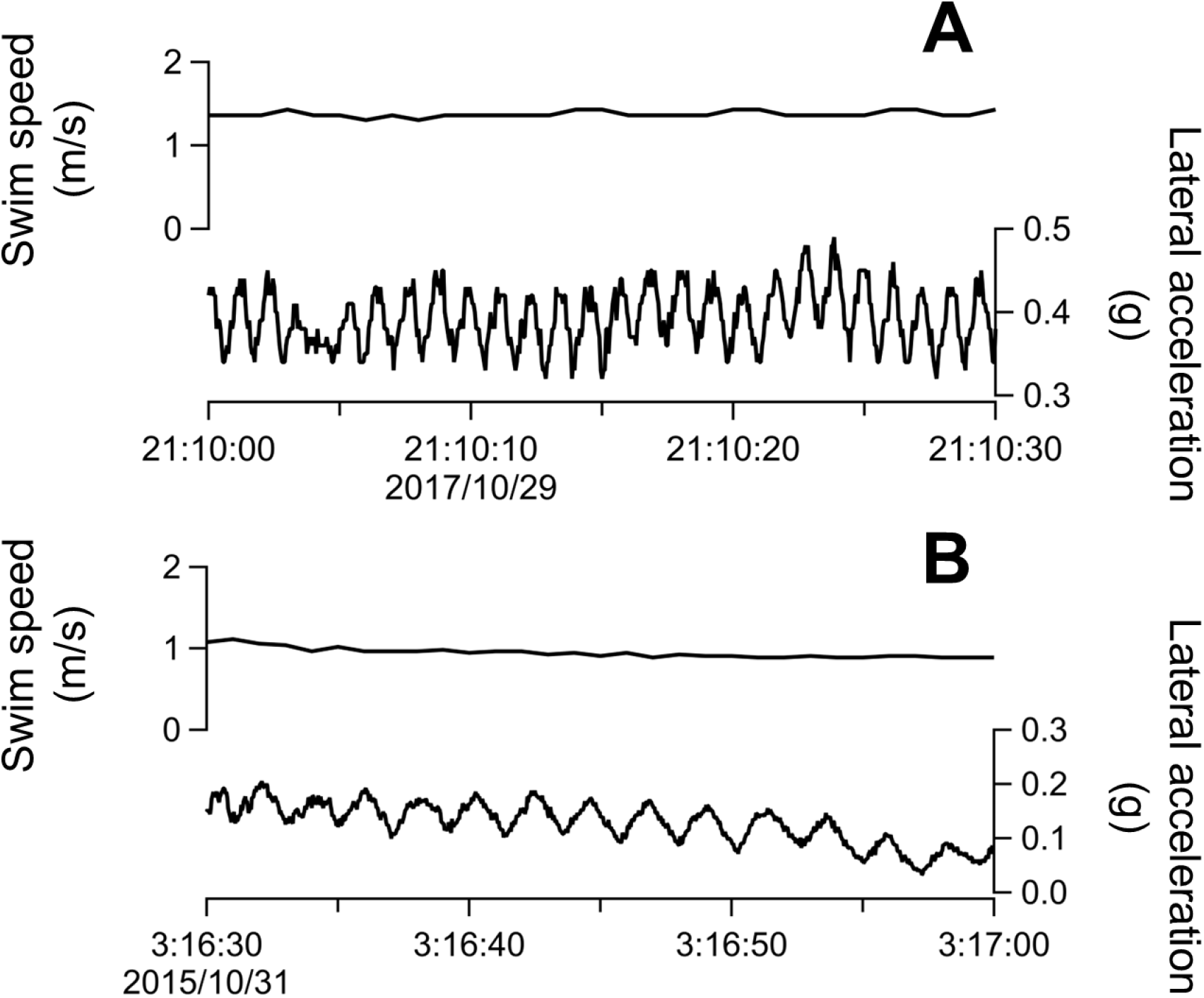
Illustrative 30-s time series of swim speed and lateral acceleration for (A) a 209 kg Atlantic bluefin tuna and (B) a 226 kg white shark. Each oscillation in lateral acceleration indicates one complete tailbeat cycle.

## DISCUSSION

### How RM-endothermic fishes cruise fast

Our comparative analyses show that the elevated cruising speeds of RM-endothermic fishes are associated primarily with increased TBF, rather than extended SL (Fig. 2A, C, E). For a given body mass and water temperature, RM-endothermic fishes exhibited 1.9 times higher cruising speed (Figs. 2A, 3A) and TBF (Figs. 2C, 3C) than ectothermic fishes, whereas SL was not significantly affected by temperature or RM endothermy (Figs. 2E, 3E). These results are consistent with the expected physiological effects of elevated RM temperature. Warmer muscles can contract more rapidly (Johnston and Brill, 1984; Rome and Sosnicki, 1990) and may therefore support higher TBF during sustained swimming. Such locomotor advantages should be particularly important in cold environments such as deep and high-latitude waters, where muscle and metabolic function are otherwise constrained (Bennett, 1984; Watanabe and Payne, 2023).

Accounting for body temperature reduced, but did not eliminate, the difference in cruising speed: RM-endothermic fishes still cruised 1.4 times faster than ectothermic fishes for a given body temperature (Fig. 3B). Thus, RM temperature elevation alone cannot fully explain the elevated cruising speed associated with RM endothermy. Other traits that support sustained high-performance swimming may also contribute, including streamlined bodies and lunate caudal fins with high aspect ratio (Bernal et al., 2001). Elevated metabolic rates in RM-endothermic fishes (Payne et al., 2026) may also increase the optimal swimming speed that minimizes cost of transport (Claireaux et al., 2006; Whitney et al., 2016). However, this result should be interpreted cautiously. Except for shortfin mako sharks, body temperatures of RM-endothermic fishes were estimated from previous studies rather than measured directly in our dataset. Moreover, the effect of RM endothermy on cruising speed was no longer statistically significant when body mass and body temperature were simultaneously included in the model (Table S4), or when whale sharks and Greenland sharks (the species occupying the extremes of body size and temperature, respectively) were excluded from the analyses (Table S5).

While RM endothermy is generally associated with higher TBF, RM-endothermic tunas and lamnid sharks retain significant differences in kinematic features. Despite similar cruising speeds (Fig. 2B), tunas showed 2.3 times higher TBF than similar-sized lamnid sharks (Figs. 2D, 4), and conversely, lamnid sharks showed 1.7 times longer SL than tunas (Fig. 2F). These results likely reflect morphological and kinematic differences between tunas and lamnid sharks. Specifically, apparently larger caudal fin area (Fig. 1) and greater tailbeat amplitude in lamnid sharks (Donley et al., 2005; Wardle et al., 1989) may allow them to generate greater thrust and travel farther with each tailbeat. In contrast, tunas appear to rely on more frequent but lower-amplitude tailbeats to achieve high-speed cruising.

Despite contrasting kinematics, both tunas and lamnid sharks may achieve similarly efficient propulsion. The Strouhal number (*St* = *fA*/*U*, where *f* is stroke frequency, *A* is stroke amplitude, and *U* is forward speed) is widely used as an indicator of efficient oscillatory propulsion, as propulsive efficiency during steady swimming is maximized when *St* is between 0.2 and 0.4 (Taylor et al., 2003). Using previously reported peak-to-peak tailbeat amplitudes (16% of FL in tunas and 26% of TL in lamnid sharks) (Donley et al., 2005; Wardle et al., 1989), *St* was estimated to be 0.23–0.45 for tunas and 0.28–0.43 for lamnid sharks in our dataset, with the majority of individuals falling within the optimal range (i.e., 0.2–0.4). Therefore, tunas and lamnid sharks may achieve near-optimal Strouhal numbers through contrasting combinations of tailbeat frequency and amplitude. Since tailbeat amplitude was not measured in our dataset, further measurements of free-swimming fish are needed to test this interpretation.

### Why tunas and lamnids evolved distinct kinematic features

The kinematic divergence between tunas and lamnid sharks raises an intriguing question: why did they evolve different solutions to achieve high-speed cruising? One possible explanation lies in skeletal type. Tunas (osteichthyans) possess bony skeletons, whereas lamnid sharks (chondrichthyans) possess cartilaginous skeletons (Currey, 2010; Porter et al., 2006). The relatively rigid skeletal structure of tunas may facilitate faster body oscillations and higher TBF, whereas the cartilaginous skeletal structure of lamnid sharks may contribute to greater tailbeat amplitudes and longer SL, although direct comparisons of their body stiffness are lacking. Therefore, the observed kinematic differences between tunas and lamnid sharks may reflect phylogenetic constraints associated with their fundamentally different skeletal types.

Differences in caudal fin aspect ratio (fin span^2^ / fin area) also provide insight into their kinematic divergence. High aspect ratios are associated with efficient cruising, whereas low aspect ratios are related to enhanced capacity for burst swimming (Webb, 1984). Lamnid sharks exhibit caudal fin aspect ratios (3–4) that exceed those of most sharks (VanderWright et al., 2024) but remain lower than those of tunas (>5) (Bernal et al., 2001) (Fig. 1A). This intermediate morphology may allow lamnid sharks to combine the capacity for long-distance migrations at high cruising speeds (Bonfil et al., 2005; Watanabe et al., 2015) with the ability to pursue or ambush highly active prey such as tunas and marine mammals (Boldrocchi et al., 2017; Martin et al., 2005; Okamura et al., 2024; Semmens et al., 2019). Consistent with this scenario, white sharks often swim at speeds lower than those that minimize cost of transport, before accelerating rapidly to ambush seals (i.e., a ‘sit-and-wait’ strategy) (Watanabe et al., 2019a; Watanabe et al., 2019b). Therefore, tunas may have specialized exclusively for sustained high-speed cruising, whereas lamnid sharks may have evolved to balance the capacity for efficient cruising with rapid acceleration. Testing this hypothesis will require more robust estimates of burst-swimming performance in free-ranging RM-endothermic fishes. Because burst-swimming events are rare and maximum speed estimates are highly sensitive to data duration and sample size, longer deployment durations and larger sample sizes will be needed.

There are several limitations to this study. First, our dataset includes only two species of bluefin tuna and three species of lamnid sharks. Broader taxonomic sampling, including both additional RM-endothermic species and closely related ectothermic species (e.g., the genus *Sarda*), would allow more robust assessments of variation among RM-endothermic fishes and of kinematic patterns associated with RM endothermy. Second, our dataset lacks direct measurements of tailbeat amplitude. Although differences in tailbeat amplitude between tunas and lamnid sharks have been reported under captive conditions (Donley et al., 2005; Wardle et al., 1989), it remains unclear whether similar patterns occur in free-ranging individuals. Reliable estimates of tailbeat amplitude in the wild are also required to estimate propulsive efficiency (Yates, 1983), which could not be assessed in this study. Future studies could address this limitation by combining biologging with video-based observations, such as drone-based video recordings from above or animal-borne video cameras oriented toward the caudal fin.

In conclusion, our comparative analyses revealed kinematic features underlying high-speed cruising in RM-endothermic fishes. RM-endothermic fishes tend to cruise faster and beat their tails more frequently than their ectothermic counterparts. Within RM-endothermic fishes, however, tunas beat their tails more frequently than lamnid sharks, while lamnid sharks travel farther with each tailbeat than tunas. These differences within RM-endothermic fishes may be partly explained by the greater caudal fin area and tailbeat amplitude in lamnid sharks, and may also reflect contrasting skeletal types, with bony skeletons in tunas (teleosts) and cartilaginous skeletons in lamnid sharks (elasmobranchs). These findings advance our knowledge of how RM-endothermic fishes cruise fast, and provide new insights into the evolution of high-performance locomotion in aquatic vertebrates.

## Acknowledgements

We sincerely thank Nobuyuki Kutsukake, Akinori Takahashi, and Hisashi Ohtsuki for their helpful comments on this study.

## Author contributions

Conceptualization: S.T., Y.Y.W.; Data curation: S.T., N.L.P., R.K., I.N., S.F., W.-C.C., J.M.S., C.G.M, Y.Y.W.; Formal analysis: S.T.; Writing-original draft: S.T.; Writing-review & editing: N.L.P., R.K., I.N., S.F., W-C.C., J.M.S., C. G. M., Y.Y.W.; Supervision: Y.Y.W.

## Data and resource availability

All relevant data and details of resources can be found within the article and its supplementary information.

