## Supplementary Information for "Comparative analyses of tailbeat frequency and stride length reveal how regionally endothermic fishes cruise fast"

**Table S1. Length–length equations used in this study.**

| Species | Equation | Reference |
|---|---|---|
| White shark *Carcharodon carcharias* | FL = 0.9442 × TL - 5.7441 | Kohler et al. (1996) |
| Salmon shark *Lamna ditropis* | TL = 1.15 × PCL + 15.19; FL = 1.08 × PCL + 6.91 | Goldman and Musick (2006) |
| Bigeye thresher shark *Alopias superciliosus* | FL = 0.5598 × TL + 17.666 | Kohler et al. (1996) |
| Whale shark *Rhincodon typus* | TL = 1.0877 × FL + 0.0865 | Meekan et al. (2020) |
| Blue shark *Prionace glauca* | FL = 0.8313 × TL + 1.3908 | Kohler et al. (1996) |
| Oceanic whitetip shark *Carcharhinus longimanus* | TL = 1.16 × FL + 7.07 | Márquez-Farías et al. (2023) |
| Greenland shark *Somniosus microcephalus* | TL = 1.0593 × FL + 4.9387 | Nielsen et al. (2014) |
| Bluntnose sixgill shark *Hexanchus griseus* | $FL = 0.476 \times TL^{1.10}$ | Mili et al. (2021) |
| Atlantic bluefin tuna *Thunnus thynnus* | TL = 1.076178 × FL + 0.55397 | Zarrad and Missaoui (2017) |
| Common dolphinfish *Coryphaena hippurus* | TL = 1.29 × FL - 9.12 | Nguyen (2024) |

TL, total length (cm); FL, fork length (cm); PCL, precaudal length (cm). For the whale shark, fish length in the equation was expressed in m, not in cm. Because no TL–FL relationship was available for Pacific bluefin tuna, the relationship reported for Atlantic bluefin tuna was used instead.

**Table S2. Length–weight equations used for body mass estimation.**

| Species | Equation | Reference |
|---|---|---|
| White shark *Carcharodon carcharias* | $W = 1.61 \times 10^{-6} \times TL^{3.309}$ | Christiansen et al. (2014) |
| Salmon shark *Lamna ditropis* | $W = 4.4 \times 10^{-5} \times PCL^{2.875}$ | Goldman and Musick (2006) |
| Shortfin mako shark *Isurus oxyrinchus* | $W = 5.2432 \times 10^{-6} \times FL^{3.1407}$ | Kohler et al. (1996) |
| Bigeye thresher shark *Alopias superciliosus* | $W = 9.1069 \times 10^{-6} \times FL^{3.0802}$ | Kohler et al. (1996) |
| Whale shark *Rhincodon typus* | $W = 3.89 \times 10^{-6} \times TL^{3.12}$ | Froese and Pauly (2026) |
| Blue shark *Prionace glauca* | $W = 3.1841 \times 10^{-6} \times FL^{3.1313}$ | Kohler et al. (1996) |
| Tiger shark *Galeocerdo cuvier* | $W = 2.5281 \times 10^{-6} \times FL^{3.2603}$ | Kohler et al. (1996) |
| Blacktip shark *Carcharhinus limbatus* | $W = 2.512 \times 10^{-9} \times TL^{3.1253}$ | Castro (1996) |
| Blacktip reef shark *Carcharhinus melanopterus* | $W = 1.004 \times 10^{-6} \times TL^{3.39}$ | Stevens (1984) |
| Grey reef shark *Carcharhinus amblyrhynchos* | $W = 1.36 \times 10^{-6} \times TL^{3.34}$ | Wetherbee et al. (1997) |
| Oceanic whitetip shark *Carcharhinus longimanus* | $W = 1.822 \times 10^{-5} \times TL^{2.78}$ | Anderson and Waheed (1990) |

| | | |
|---|---|---|
| Greenland shark<br>*Somniosus microcephalus* | $W = 4.416 \times 10^{-6} \times TL^{3.1346}$ | Nielsen et al. (2014) |
| Bluntnose sixgill shark<br>*Hexanchus griseus* | $W = -37.5 + 0.0664 \times TL - 3.11 \times 10^{-5} \times TL^{2} + 10^{-8} \times TL^{3}$ | Ebert (1984) |
| Broadnose sevengill shark<br>*Notorynchus cepedianus* | $W = -3.39 + 0.0123 \times TL - 1.58 \times 10^{-5} \times TL^{2} + 1.01 \times 10^{-8} \times TL^{3}$ | Ebert (1984) |
| Atlantic bluefin tuna<br>*Thunnus thynnus* | $W = 2 \times 10^{-5} \times TL^{2.96}$ | Sinovčić et al. (2004) |
| Pacific bluefin tuna<br>*Thunnus orientalis* | $W = 1.7117 \times 10^{-5} \times FL^{3.0382}$ | Kai (2007) |
| Common dolphinfish<br>*Coryphaena hippurus* | $W = 1.0693 \times 10^{-5} \times FL^{2.9337}$ | Uchiyama and Boggs (2006) |
| Striped marlin<br>*Kajikia audax* | $W = 4.68 \times 10^{-6} \times EFL^{3.16}$ | Sun et al. (2011) |
| Greater amberjack<br>*Seriola dumerili* | $W = 1.9 \times 10^{-5} \times TL^{2.8726}$ | Mohamed et al. (2018) |
| Yellowtail amberjack<br>*Seriola lalandi* | $W = 3.515 \times 10^{-8} \times FL^{2.845}$ | Taylor and Willis (1998) |

W, body mass (kg); TL, total length (cm); FL, fork length (cm); PCL, precaudal length (cm); EFL, eye-fork length (cm). For the blacktip shark, bluntnose sixgill shark, broadnose sevengill shark, and yellowtail amberjack, fish length in the equation is expressed in mm rather than cm. For the salmon shark, PCL was estimated from TL using a length–length relationship (Table S1; Goldman and Musick, 2006), then used to estimate body mass. Similarly, for oceanic whitetip sharks, TL was estimated from FL using a length–length relationship (Table S1; Márquez-Farías et al., 2023), then used to estimate body mass. For the striped marlin, TL was measured and body mass was estimated at capture. Because EFL was not measured and no published TL–EFL relationship was available for this species, EFL was estimated from body mass using the length–weight relationship reported by Sun et al. (2011).

**Table S3. Body size and kinematic parameters for 66 individuals from 20 fish species.**

| Species | Total length (m) | Fork length (m) | Body mass (kg)[b] | Analyzed Duration (h)[c] | Cruising speed (m/s) | Tailbeat frequency (Hz) | Stride length (m) | Water temp. (℃) | Body temp. (℃) |
|---|---|---|---|---|---|---|---|---|---|
| *Carcharodon carcharias* | 3.30 | 3.06[a] | 347 | 12.0 | 0.87 | 0.39 | 2.23 | 15.5 | 26.4 |
| *Carcharodon carcharias* | 3.20 | 2.96[a] | 314 | 14.7 | 0.83 | 0.37 | 2.24 | 15.4 | 26.4 |
| *Carcharodon carcharias* | 4.20 | 3.91[a] | 771 | 13.5 | 0.86 | 0.27 | 3.19 | 15.1 | 26.4 |
| *Carcharodon carcharias* | 4.30 | 4.00[a] | 834 | 26.5 | 0.91 | 0.27 | 3.37 | 15.2 | 26.4 |
| *Carcharodon carcharias* | 3.80 | 3.53[a] | 554 | 17.0 | 1.37 | 0.47 | 2.91 | 15.5 | 26.4 |
| *Carcharodon carcharias* | 3.70 | 3.44[a] | 507 | 7.9 | 1.07 | 0.47 | 2.28 | 15.4 | 26.4 |
| *Carcharodon carcharias* | 2.90 | 2.68[a] | 226 | 8.7 | 0.80 | 0.38 | 2.11 | 15.5 | 26.4 |
| *Lamna ditropis* | 2.15 | 1.95[a] | 121 | 4.1 | 1.12 | 0.56 | 2.00 | 13.4 | 25.3 |
| *Isurus oxyrinchus* | 1.70 | 1.63 | 46.5 | 12.3 | 0.59 | 0.57 | 1.04 | 20.7 | 22.4 |
| *Isurus oxyrinchus* | 1.45 | 1.29 | 22.3 | 22.4 | 0.58 | 0.60 | 0.97 | 20.0 | 21.7 |
| *Isurus oxyrinchus* | 1.70 | 1.59 | 43.0 | 18.2 | 0.63 | 0.59 | 1.07 | 18.8 | 23.0 |
| *Alopias superciliosus* | 2.70 | 1.69[a] | 66.3 | 14.6 | 0.42 | 0.35 | 1.20 | 15.7 | 15.7 |
| *Rhincodon typus* | 6.94 | 6.30[a] | 2850 | 125 | 0.46 | 0.22 | 2.09 | 26.7 | 26.7 |
| *Rhincodon typus* | 4.44 | 4.00[a] | 708 | 12.5 | 0.57 | 0.24 | 2.38 | 23.4 | 23.4 |
| *Rhincodon typus* | 7.24 | 6.58[a] | 3250 | 64.1 | 0.48 | 0.21 | 2.29 | 27.4 | 27.4 |
| *Prionace glauca* | 1.92[a] | 1.61 | 25.9 | 9.5 | 0.38 | 0.51 | 0.75 | 20.4 | 20.4 |
| *Prionace glauca* | 2.08[a] | 1.74 | 33.0 | 7.7 | 0.38 | 0.37 | 1.03 | 19.8 | 19.8 |
| *Galeocerdo cuvier* | 3.87 | 3.22 | 379 | 3.3 | 0.93 | 0.35 | 2.66 | 24.2 | 24.2 |
| *Galeocerdo cuvier* | 3.80 | 3.05 | 318 | 138 | 0.83 | 0.35 | 2.37 | 26.0 | 26.0 |
| *Galeocerdo cuvier* | 3.43 | 2.89 | 267 | 3.7 | 0.99 | 0.34 | 2.91 | 25.2 | 25.2 |
| *Galeocerdo cuvier* | 2.43 | 1.96 | 75.2 | 96.8 | 0.74 | 0.42 | 1.76 | 25.0 | 25.0 |
| *Galeocerdo cuvier* | 3.61 | 2.85 | 255 | 14.5 | 1.03 | 0.37 | 2.78 | 25.0 | 25.0 |
| *Galeocerdo cuvier* | 2.47 | 1.97 | 76.5 | 19.1 | 0.86 | 0.48 | 1.79 | 25.2 | 25.2 |
| *Galeocerdo cuvier* | 3.92 | 3.28 | 403 | 122 | 0.58 | 0.32 | 1.81 | 23.8 | 23.8 |
| *Galeocerdo cuvier* | 2.76 | 2.20 | 110 | 166 | 0.58 | 0.39 | 1.49 | 23.6 | 23.6 |
| *Galeocerdo cuvier* | 2.24 | 1.79 | 55.9 | 9.2 | 0.72 | 0.47 | 1.53 | 23.7 | 23.7 |
| *Galeocerdo cuvier* | 1.90 | 1.55 | 35.0 | 12.5 | 0.68 | 0.61 | 1.11 | 23.7 | 23.7 |
| *Carcharhinus limbatus* | 1.46 | 1.18 | 19.5 | 63.8 | 0.47 | 0.83 | 0.57 | 23.9 | 23.9 |
| *Carcharhinus melanopterus* | 1.13 | 0.93 | 9.2 | 59.5 | 0.51 | 0.80 | 0.64 | 28.4 | 28.4 |
| *Carcharhinus melanopterus* | 1.27 | 1.05 | 13.6 | 24.5 | 0.52 | 0.83 | 0.63 | 29.0 | 29.0 |
| *Carcharhinus amblyrhynchos* | 1.58 | 1.29 | 30.0 | 63.2 | 0.54 | 0.49 | 1.10 | 27.2 | 27.2 |
| *Carcharhinus longimanus* | 2.58[a] | 2.16 | 92.2 | 73.1 | 0.67 | 0.39 | 1.72 | 25.5 | 25.5 |
| *Carcharhinus longimanus* | 2.09[a] | 1.74 | 51.4 | 9.4 | 0.73 | 0.47 | 1.55 | 25.7 | 25.7 |
| *Carcharhinus longimanus* | 2.80[a] | 2.35 | 116 | 28.5 | 0.75 | 0.37 | 2.03 | 26.0 | 26.0 |

| | | | | | | | | | |
|---|---|---|---|---|---|---|---|---|---|
| *Carcharhinus longimanus* | 2.51[a] | 2.10 | 85.4 | 35.8 | 0.63 | 0.40 | 1.58 | 25.2 | 25.2 |
| *Somniosus microcephalus* | 2.56 | 2.37[a] | 156 | 8.3 | 0.19 | 0.14 | 1.36 | 1.0 | 1.0 |
| *Somniosus microcephalus* | 3.00 | 2.87 | 257 | 8.6 | 0.26 | 0.16 | 1.63 | 1.0 | 1.0 |
| *Somniosus microcephalus* | 3.30 | 3.10 | 346 | 19.3 | 0.26 | 0.14 | 1.86 | 0.7 | 0.7 |
| *Somniosus microcephalus* | 2.22 | 2.15 | 100 | 9.3 | 0.22 | 0.14 | 1.57 | 0.3 | 0.3 |
| *Somniosus microcephalus* | 3.00 | 2.80 | 257 | 48.9 | 0.23 | 0.14 | 1.64 | 0.7 | 0.7 |
| *Somniosus microcephalus* | 2.86 | 2.65[a] | 221 | 12.1 | 0.34 | 0.18 | 1.89 | 0.9 | 0.9 |
| *Somniosus microcephalus* | 2.23 | 2.06[a] | 101 | 13.3 | 0.19 | 0.18 | 1.06 | 1.0 | 1.0 |
| *Hexanchus griseus* | 4.61 | 4.05[a] | 587 | 104 | 0.45 | 0.14 | 3.21 | 10.6 | 10.6 |
| *Hexanchus griseus* | 4.16 | 3.62[a] | 420 | 13.4 | 0.51 | 0.22 | 2.32 | 12.1 | 12.1 |
| *Hexanchus griseus* | 3.33 | 2.83[a] | 208 | 112 | 0.35 | 0.18 | 1.94 | 9.5 | 9.5 |
| *Hexanchus griseus* | 4.40 | 3.85[a] | 504 | 49.2 | 0.30 | 0.17 | 1.76 | 9.5 | 9.5 |
| *Notorynchus cepedianus* | 1.95 | 1.52 | 35.4 | 77.7 | 0.45 | 0.42 | 1.07 | 17.0 | 17.0 |
| *Notorynchus cepedianus* | 2.10 | 1.63 | 46.3 | 8.8 | 0.43 | 0.39 | 1.10 | 17.2 | 17.2 |
| *Notorynchus cepedianus* | 2.50 | 1.93 | 86.4 | 21.2 | 0.47 | 0.49 | 0.96 | 17.0 | 17.0 |
| *Notorynchus cepedianus* | 1.88 | 1.46 | 31.0 | 65.5 | 0.47 | 0.43 | 1.09 | 17.3 | 17.3 |
| *Thunnus thynnus* | 2.35 | 2.18[a] | 209 | 1.3 | 1.46 | 0.95 | 1.54 | 12.8 | 24.3 |
| *Thunnus thynnus* | 2.05 | 1.90[a] | 139 | 13.8 | 0.87 | 0.96 | 0.91 | 13.4 | 24.6 |
| *Thunnus thynnus* | 2.35 | 2.18[a] | 209 | 1.4 | 0.98 | 0.76 | 1.29 | 13.2 | 24.5 |
| *Thunnus thynnus* | 2.20 | 2.04[a] | 172 | 6.9 | 0.96 | 0.85 | 1.13 | 13.0 | 24.4 |
| *Thunnus orientalis* | 1.59[a] | 1.47 | 65.8 | 5.3 | 1.44 | 1.53 | 0.94 | 24.1 | 30.9 |
| *Thunnus orientalis* | 0.83[a] | 0.77 | 9.2 | 1.4 | 0.78 | 2.28 | 0.34 | 15.9 | 19.4 |
| *Thunnus orientalis* | 0.92[a] | 0.85 | 12.5 | 32.1 | 1.07 | 2.49 | 0.43 | 17.1 | 21.0 |
| *Thunnus orientalis* | 0.95[a] | 0.88 | 13.8 | 9.3 | 0.79 | 2.50 | 0.32 | 19.1 | 23.1 |
| *Coryphaena hippurus* | 1.01[a] | 0.85 | 4.9 | 38.7 | 0.74 | 0.88 | 0.84 | 19.9 | 19.9 |
| *Coryphaena hippurus* | 0.93[a] | 0.79 | 3.9 | 11.3 | 0.84 | 1.85 | 0.45 | 27.5 | 27.5 |
| *Coryphaena hippurus* | 0.97[a] | 0.82 | 4.4 | 12.0 | 0.84 | 1.48 | 0.57 | 24.7 | 24.7 |
| *Coryphaena hippurus* | 1.07[a] | 0.90 | 5.8 | 29.8 | 0.68 | 0.94 | 0.72 | 24.5 | 24.5 |
| *Coryphaena hippurus* | 0.98[a] | 0.83 | 4.6 | 17.8 | 0.50 | 1.29 | 0.39 | 22.9 | 22.9 |
| *Kajikia audax* | 2.50 | 1.68[a] | 50 | 7.9 | 0.69 | 0.71 | 0.97 | 22.7 | 22.7 |
| *Seriola dumerili* | 0.85 | 0.81 | 6.6 | 5.2 | 1.08 | 1.76 | 0.61 | 20.1 | 20.1 |
| *Seriola lalandi* | 1.04 | 0.95 | 10.4 | 30.3 | 0.59 | 1.09 | 0.54 | 21.7 | 21.7 |

[a] Missing TL or FL values were estimated using species-specific TL–FL relationships (Table S1), except for the striped marlin, for which eye-fork length (EFL) was estimated from body mass using the length–weight relationship (Table S2).

[b] Estimated from length–weight relationships (Table S2), except for the striped marlin, for which body mass was estimated at capture.

[c] Duration excluding the first 6 h after capture and including only periods of horizontal swimming (see Materials and Methods).

**Table S4. Posterior parameter estimates for full models from the MCMCglmm analyses.**

| Comparison | Model | Predictor | Posterior Mean | 95% credible interval | |
|---|---|---|---|---|---|
| | | | | Lower | Upper |
| RM endothermy vs. Ectothermy | $\log_{10}$(Speed) ~ $\log_{10}$(Mass) + Water temp + Endothermy | $\log_{10}$(Mass) | 0.067 | -0.0023 | 0.13 |
| | | Water temp | 0.015 | 0.0064 | 0.023 |
| | | Endothermy | 0.26 | 0.11 | 0.42 |
| | $\log_{10}$(Speed) ~ $\log_{10}$(Mass) + Body temp + Endothermy | $\log_{10}$(Mass) | 0.044 | -0.017 | 0.099 |
| | | Body temp | 0.017 | 0.010 | 0.025 |
| | | Endothermy | 0.14 | -0.0088 | 0.28 |
| | $\log_{10}$(TBF) ~ $\log_{10}$(Mass) + Water temp + Endothermy | $\log_{10}$(Mass) | -0.23 | -0.29 | -0.18 |
| | | Water temp | 0.015 | 0.0081 | 0.023 |
| | | Endothermy | 0.29 | 0.14 | 0.44 |
| | $\log_{10}$(TBF) ~ $\log_{10}$(Mass) + Body temp + Endothermy | $\log_{10}$(Mass) | -0.26 | -0.30 | -0.20 |
| | | Body temp | 0.014 | 0.0073 | 0.021 |
| | | Endothermy | 0.17 | 0.037 | 0.33 |
| | $\log_{10}$(SL) ~ $\log_{10}$(Mass) + Water temp + Endothermy | $\log_{10}$(Mass) | 0.31 | 0.24 | 0.37 |
| | | Water temp | -0.00087 | -0.0095 | 0.0070 |
| | | Endothermy | -0.036 | -0.20 | 0.10 |
| | $\log_{10}$(SL) ~ $\log_{10}$(Mass) + Body temp + Endothermy | $\log_{10}$(Mass) | 0.31 | 0.25 | 0.37 |
| | | Body temp | 0.0022 | -0.0062 | 0.010 |
| | | Endothermy | -0.041 | -0.20 | 0.11 |

| | | | | | |
|---|---|---|---|---|---|
| Tuna vs. Lamnid shark | $\log_{10}$(Speed) ~ $\log_{10}$(Mass) + Water temp + Clade | $\log_{10}$(Mass) | 0.18 | 0.012 | 0.33 |
| | | Water temp | 0.0054 | -0.017 | 0.027 |
| | | Clade | 0.16 | -0.24 | 0.48 |
| | $\log_{10}$(Speed) ~ $\log_{10}$(Mass) + Body temp + Clade | $\log_{10}$(Mass) | 0.11 | -0.063 | 0.33 |
| | | Body temp | 0.013 | -0.011 | 0.039 |
| | | Clade | 0.14 | -0.12 | 0.44 |
| | $\log_{10}$(TBF) ~ $\log_{10}$(Mass) + Water temp + Clade | $\log_{10}$(Mass) | -0.25 | -0.37 | -0.10 |
| | | Water temp | 0.0023 | -0.017 | 0.022 |
| | | Clade | 0.36 | 0.093 | 0.65 |
| | $\log_{10}$(TBF) ~ $\log_{10}$(Mass) + Body temp + Clade | $\log_{10}$(Mass) | -0.27 | -0.43 | -0.090 |
| | | Body temp | 0.0041 | -0.020 | 0.026 |
| | | Clade | 0.36 | 0.093 | 0.60 |
| | $\log_{10}$(SL) ~ $\log_{10}$(Mass) + Water temp + Clade | $\log_{10}$(Mass) | 0.42 | 0.31 | 0.55 |
| | | Water temp | 0.0038 | -0.012 | 0.021 |
| | | Clade | -0.22 | -0.43 | -0.013 |
| | $\log_{10}$(SL) ~ $\log_{10}$(Mass) + Body temp + Clade | $\log_{10}$(Mass) | 0.38 | 0.25 | 0.53 |
| | | Body temp | 0.0079 | -0.0097 | 0.025 |
| | | Clade | -0.23 | -0.41 | -0.056 |

Water and body temperature were included separately to avoid multicollinearity. Effects of predictors were considered statistically significant when the 95% credible intervals did not include zero.

**Table S5. Posterior parameter estimates for final models from the MCMCglmm analyses excluding whale sharks and Greenland sharks.**

| Model | Predictor | Posterior Mean | 95% credible interval | |
|---|---|---|---|---|
| | | | Lower | Upper |
| $\log_{10}$(Speed) ~ $\log_{10}$(Mass) + Water temp + Endothermy | $\log_{10}$(Mass) | 0.097 | 0.033 | 0.17 |
| | Water temp | 0.016 | 0.0056 | 0.027 |
| | Endothermy | 0.23 | 0.069 | 0.38 |
| $\log_{10}$(Speed) ~ $\log_{10}$(Mass) + Body temp | $\log_{10}$(Mass) | 0.076 | 0.0090 | 0.14 |
| | Body temp | 0.020 | 0.011 | 0.029 |
| $\log_{10}$(TBF) ~ $\log_{10}$(Mass) + Water temp + Endothermy | $\log_{10}$(Mass) | -0.25 | -0.30 | -0.18 |
| | Water temp | 0.013 | 0.0038 | 0.024 |
| | Endothermy | 0.28 | 0.11 | 0.43 |
| $\log_{10}$(TBF) ~ $\log_{10}$(Mass) + Body temp + Endothermy | $\log_{10}$(Mass) | -0.27 | -0.33 | -0.20 |
| | Body temp | 0.012 | 0.0028 | 0.021 |
| | Endothermy | 0.19 | 0.051 | 0.36 |
| $\log_{10}$(SL) ~ $\log_{10}$(Mass) | $\log_{10}$(Mass) | 0.34 | 0.28 | 0.41 |

Water and body temperature were included separately to avoid multicollinearity. Final models were obtained by sequentially removing predictors with 95% credible intervals overlapping zero.

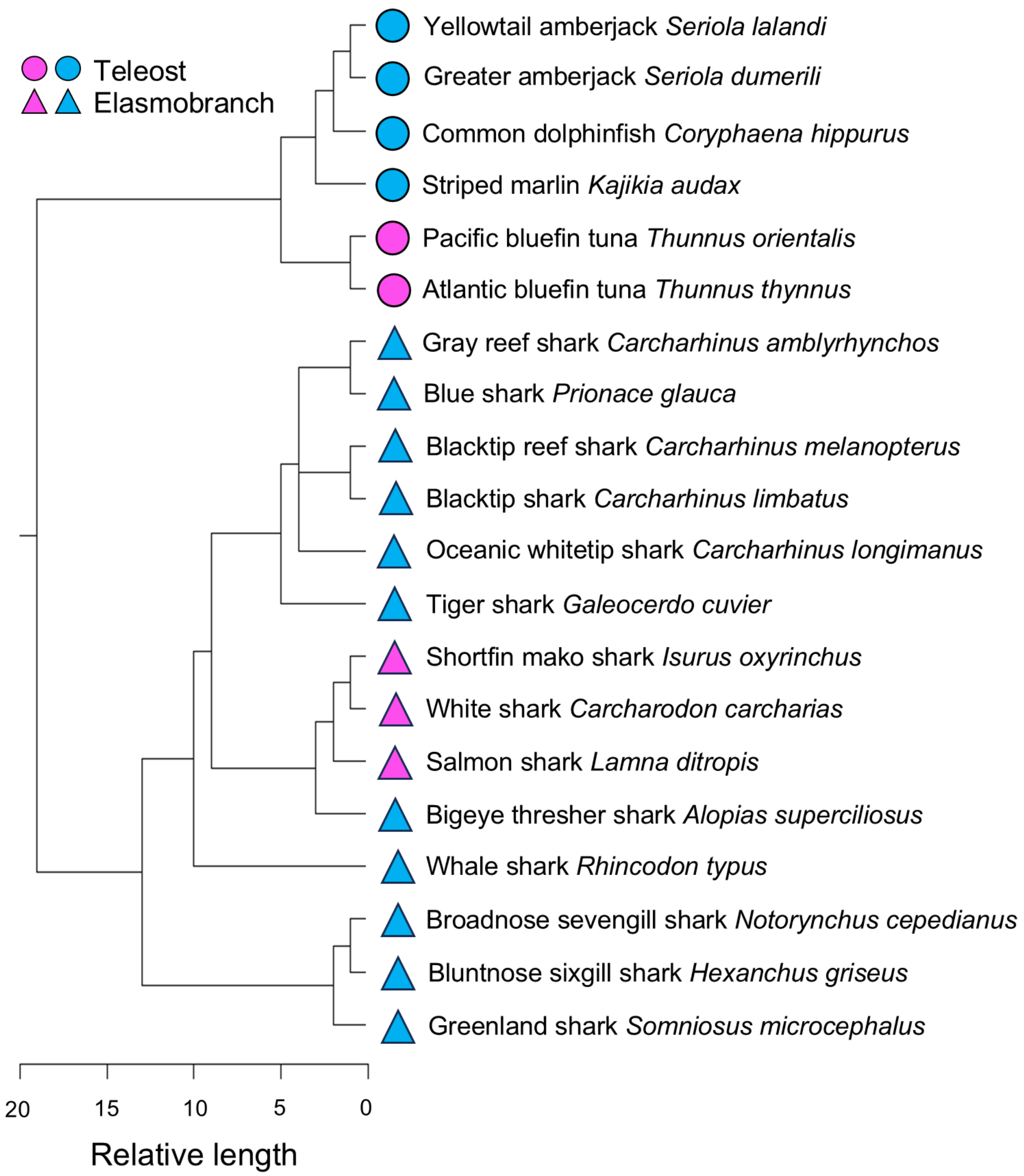


**Figure S1. Phylogenetic tree used for MCMCglmm analyses.** Marker color indicates thermal strategy (pink, RM-endothermy; sky blue, ectothermy), and marker shape indicates taxonomic group (triangles, elasmobranchs; circles, teleosts).
